# FlowAgent: A Modular Agent-Based System for Automated Workflow Management and Data Interpretation

**DOI:** 10.1101/2025.03.06.641728

**Authors:** Joshua Philpott, Alina Kurjan, Adam P Cribbs

**Affiliations:** Entelo Bio, BioEscalator, Innovation Building, University of Oxford, Old Road Campus, Roosevelt Drive, Oxford, OX3 7FZ, UK

## Abstract

Reproducibility, automation, and flexibility remain persistent challenges in bioinformatics, where complex workflows require integration of diverse computational tools, rigorous quality control, and dynamic adaptability to evolving datasets. Existing workflow managers often trade off usability for flexibility, limiting accessibility for non-specialists and enforcing rigid execution pipelines. We present FlowAgent, an adaptive agent-based framework that embeds computational heuristics and intelligent decision-making into workflow management. Unlike static pipeline managers, FlowAgent leverages agentic systems for context-aware automation, enhancing error detection, correction, and real-time optimisation while minimising manual intervention. By integrating bioinformatics tools with adaptive control, FlowAgent ensures robust quality control and enables workflow execution through an intuitive natural language interface. This approach democratises advanced computational methods for biologists while allowing bioinformaticians to focus on high-level analysis. FlowAgent redefines workflow automation, making bioinformatics pipelines more intelligent, efficient, and accessible, ultimately accelerating scientific discovery across diverse omics data modalities.

**Availability and implementation:** FlowAgent is available with GLPv3 license at https://github.com/EnteloBio/flowagent

## Introduction

Bioinformatics is a rapidly evolving field that applies computational methods to analyse and interpret complex biological data, driving advances in genomics, transcriptomics, and multi-omics research. The rise of high-throughput sequencing technologies, including single-cell RNA sequencing, spatial transcriptomics, and 3D genome analysis, has exponentially increased data volume and complexity, necessitating sophisticated computational workflows capable of integrating challenging analytical steps such as data preprocessing, quality control, and statistical interpretation(Shade and Teal 2015).

Traditional workflow management systems (WMS) (Reiter, Brooks et al. 2021), like Nextflow (Di Tommaso, Chatzou et al. 2017), Snakemake (Koster and Rahmann 2012), CGAT-core (Cribbs, Luna-Valero et al. 2019), and Galaxy Workflow Manager (Galaxy 2024) provide structured frameworks for reproducible analysis. However, these systems depend on static, script-based configurations that demand substantial programming expertise, limiting accessibility, scalability, and real-time adaptability to evolving analytical challenges. Although Galaxy’s web-based interface mitigates scripting requirements, it still lacks real-time adaptability and AI-driven workflow automation.

Recent advances in large language models (LLMs) have enabled agentic systems that automate workflow design, task execution, and quality control. Tools such as AutoBA (Zhou, Zhang et al. 2023), ChemCrow (A, Cox et al. 2024), and CellAgent (Xiao, Liu et al. 2024) demonstrate the potential of multi-agent architectures for automated computational analyses. However, these early systems face critical limitations in scalability, error detection, and adaptive workflow optimisation. Despite improvements using role-based agents, Retrieval-Augmented Generation, and memory management strategies to improve task coordination and pipeline consistency (Su, Long et al. 2025), current approaches remain predominantly task-driven and fall short of providing real-time quality control, high-performance computing (HPC) and cloud environment integration, context-aware decision-making, and fully adaptive workflow generation essential for large-scale bioinformatics analyses.

### FlowAgent: an adaptive agent-based workflow solution

To address these limitations, we introduce FlowAgent, an adaptive agent-based workflow management framework that redefines the design, execution, and optimisation of bioinformatics workflows. FlowAgent eliminates scripting barriers of traditional WMS by using LLMs for natural language workflow generation, allowing researchers to describe workflows in plain English and have them translated into structured bioinformatics pipelines. Moreover, the framework incorporates intelligent quality control by deploying agents that actively monitor tool version compatibility, dataset quality, and computational performance, providing real-time recommendations to improve accuracy. It also streamlines deployment automatically by installing any required software that is not already available in the environment.

In addition to these features, FlowAgent dynamically adapts workflow execution using context-aware decision-making to optimise parameter selection based on experimental objectives, tool selection, and dataset characteristics. The framework ensures version intelligence by actively tracking and validating software dependencies, thereby preventing disruptive version mismatches. Comprehensive, automated reporting further distinguishes FlowAgent by delivering actionable insights, performance metrics, and tool-specific recommendations beyond traditional execution logs.

By integrating LLM-based workflow design, adaptive execution, and intelligent error correction, FlowAgent transcends the limitations of static pipelines to offer a self-optimising, user-friendly, and highly scalable solution. Unlike existing WMS and agent-based tools, FlowAgent not only automates workflow management but also actively refines execution based on computational performance. This ensures that both experimental biologists and computational scientists can conduct high-performance bioinformatics analyses without requiring extensive coding expertise.

## Methods

### Design of FlowAgent

FlowAgent is an intelligent workflow management system that automates bioinformatics analyses using a structured, agent-based framework. It operates in three stages: workflow planning, adaptive execution, and result generation with intelligent reporting (Figure 1).

**Figure 1:**
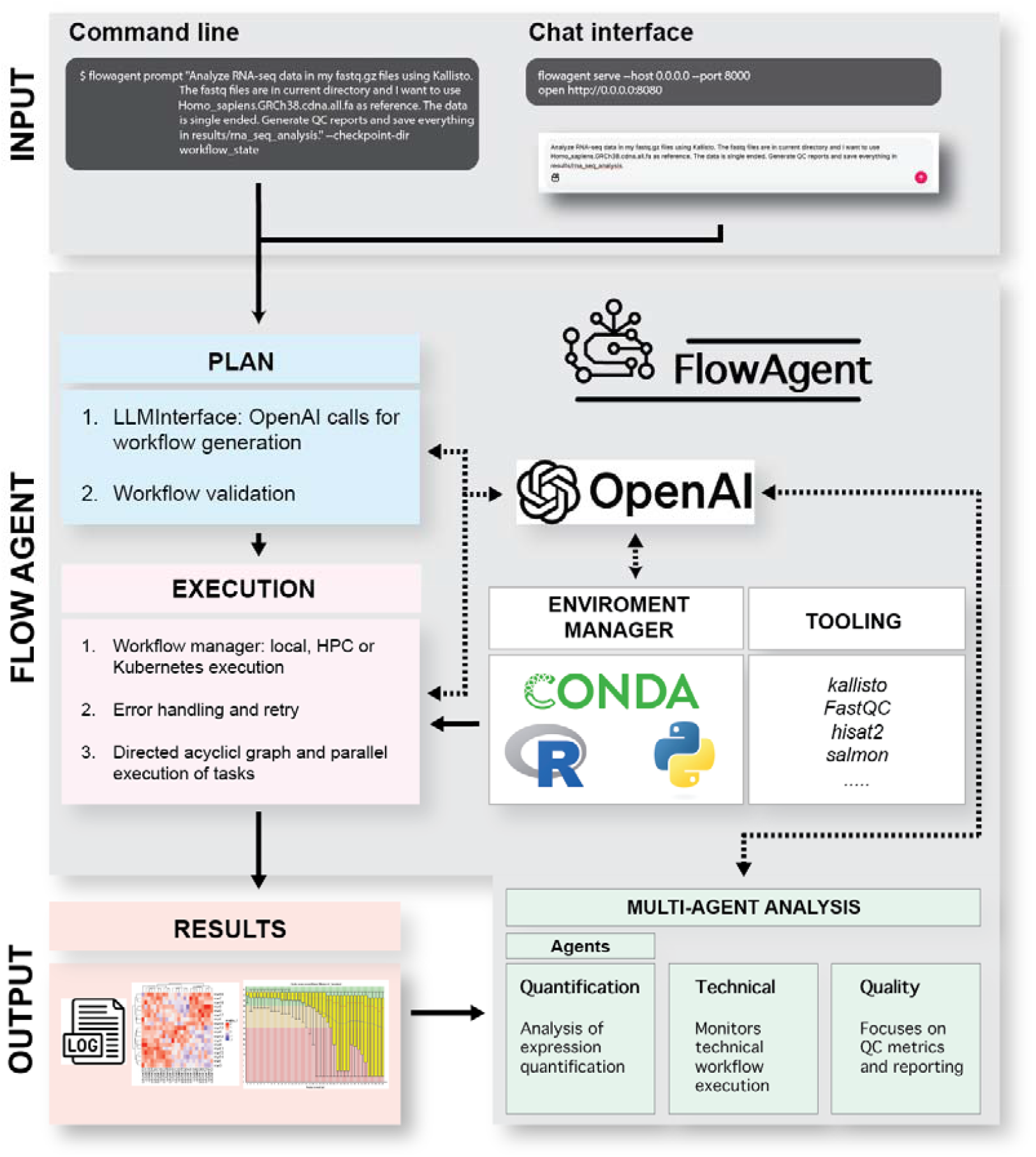
Design of FlowAgent. FlowAgent is an agentic workflow management system designed to automate, validate, and execute computational workflows across multiple domains. The system receives user input via a command-line interface or a chat-based interface, where users specify tasks in natural language. In the planning phase, FlowAgent leverages LLM-based workflow generation and validation to construct executable workflows. The execution phase orchestrates computational workflows across local, HPC, or Kubernetes environments, incorporating error handling, retry mechanisms, and real-time agentic reporting. The environment manager ensures seamless integration of dependencies using Conda, R, and Python, while the tooling layer supports various software, such as Kallisto, FastQC, Hisat2, and Salmon, among others. The system generates structured results, including logs and visual reports, which are processed through a multi-agent analysis framework. Dedicated agents assess quantification, technical workflow execution, and quality control metrics, ensuring robust, reproducible, and interpretable computational analyses across diverse applications.

In the planning phase, users provide natural language descriptions of their desired analyses via a command line or chat-based interface. Through its API-driven integration with OpenAI models and intelligent automation, FlowAgent translates these descriptions into structured workflows. For example, a request like “I want to download data from GEO with the accession number GSE186412. I then want to analyse the bulk RNA-seq data using kallisto with the reference Mus_musculus.GRCm39.cdna.all.fa. The data is paired ended” is automatically processed to extract the experimental context, desired outcomes, and analysis requirements. FlowAgent subsequently constructs an executable workflow by identifying necessary analytical steps, selecting optimal computational tools, configuring parameters, and mapping dependencies into a directed acyclic graph (DAG) that represents the entire analysis pipeline (Figure 2). The planning phase can be further refined by modifying user prompts to specify additional constraints, such as computational resource limits, alternative analytical approaches, or quality control parameters, ensuring greater adaptability to diverse research needs.

**Figure 2:**
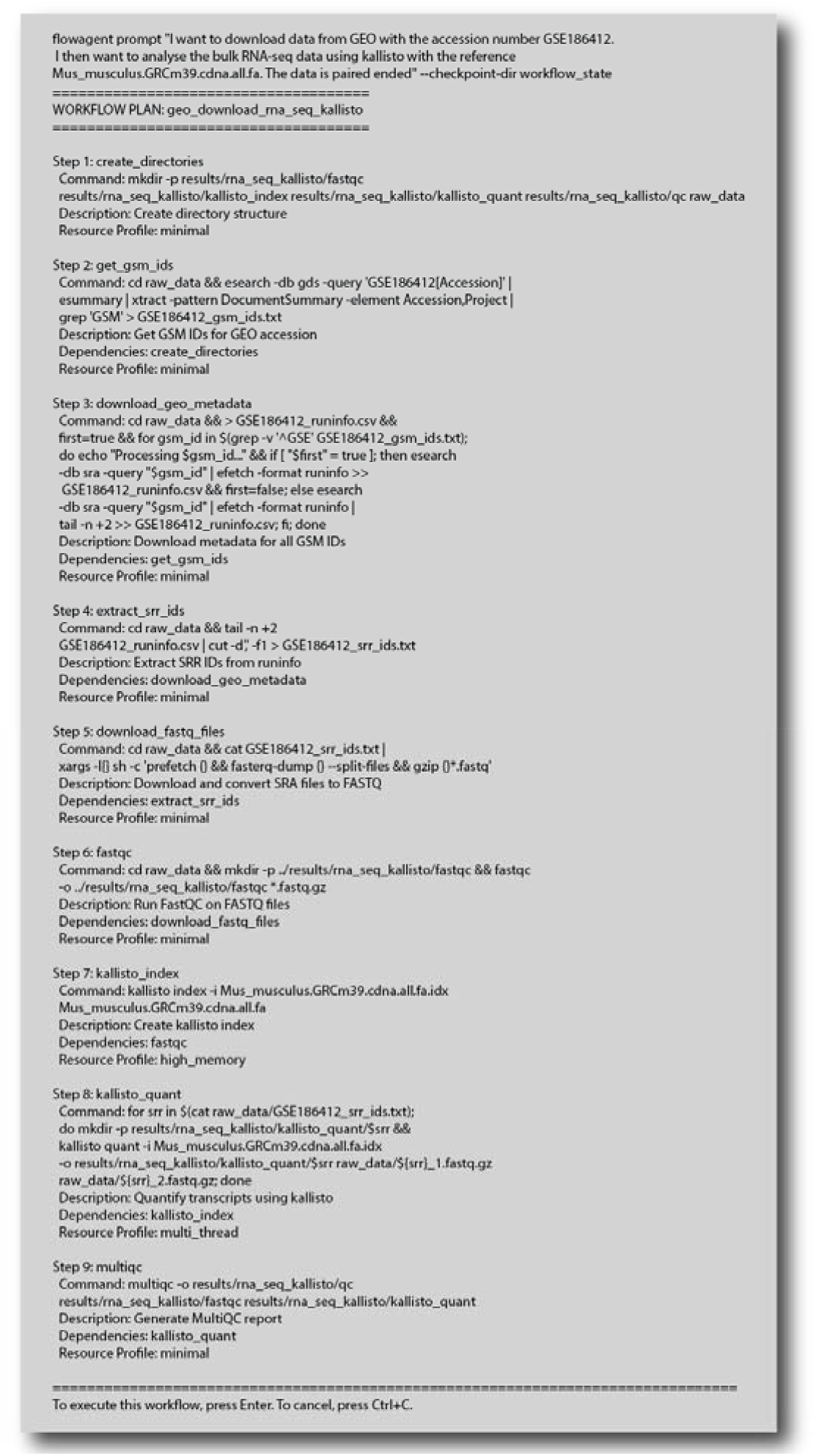
Automated workflow planning using FlowAgent. Initiated via a natural language prompt, FlowAgent constructs a structured workflow plan that identifies the required analysis steps, selects optimal computational tools, and configures command parameters, presenting a DAG in written form. When executed, this workflow: (1) creates directories, (2) accesses GSM IDs, (3) extracts associated metadata, (4) extracts SRR IDs, (5) downloads bulk RNA-seq data FASTQ files from the Gene Expression Omnibus (GEO) (Accession: GSE186412), (6) performs quality control using FastQC, (7) indexes the transcriptome with kallisto, (8) quantifies the bulk RNA-seq data transcripts, and (9) performs final quality assessment with MultiQC.

Following workflow execution, FlowAgent processes data according to the defined steps while dynamically refining analyses with an intelligent, agent-based reporting system. This system comprises three specialised agents that work collaboratively to ensure robust, high-quality outcomes (Figure 1). These agents oversee overall pipeline performance by tracking software versions, resource utilisation, and computational efficiency, identifying anomalies that could compromise execution. Together, these agents handle workflow validation, resource allocation, environment setup, and package dependency resolution, providing real-time workflow parameter optimisation and adaptive corrections that prevent cascading errors and maintain the integrity of bioinformatics analyses. Thus, FlowAgent ensures contextual adaptation by dynamically refining workflows based on experimental conditions, incorporating best practices, and handling technical requirements such as quality control thresholds.

To maximise computational flexibility and scalability, FlowAgent supports execution across HPC clusters, cloud environments, and Kubernetes-based orchestration systems. This ensures efficient resource management and adaptability to different infrastructure requirements, allowing users to leverage distributed computing for large-scale analyses while maintaining reproducibility. The environment manager ensures seamless integration of dependencies using Conda, R, and Python, while the tooling layer supports various bioinformatics software, including Kallisto, FastQC, Hisat2, and Salmon, among others. Through this modular and infrastructure-agnostic design, FlowAgent optimises computational performance across diverse research environments.

## Results

### Benchmarking API models

To assess the performance of FlowAgent across different OpenAI models and determine the optimal model for execution, we conducted a benchmarking study comparing gpt-3.5-turbo, gpt-4, gpt-4-turbo, gpt-4.5-preview, and gpt-4o (Table 1). The evaluation considered syntax correctness, execution correctness, response time, token usage, and cost efficiency. gpt-3.5-turbo exhibited the lowest syntax and execution correctness (40%), while all gpt-4-based models achieved 100% correctness. gpt-4o demonstrated the fastest execution time (4.89 min), outperforming gpt-4.5-preview (7.14 min), though gpt-4-turbo required the most API calls (10). In terms of cost, gpt-4 was the most expensive ($0.346), while gpt-4-turbo incurred a cost of $0.16. Report detail also varied, with gpt-4.5-preview generating the highest average number of reporting lines (71), compared to gpt-3.5-turbo (9). These findings provide insight into the trade-offs between computational efficiency, response accuracy, and cost, informing the optimal model selection for FlowAgent.

**Table 1:**
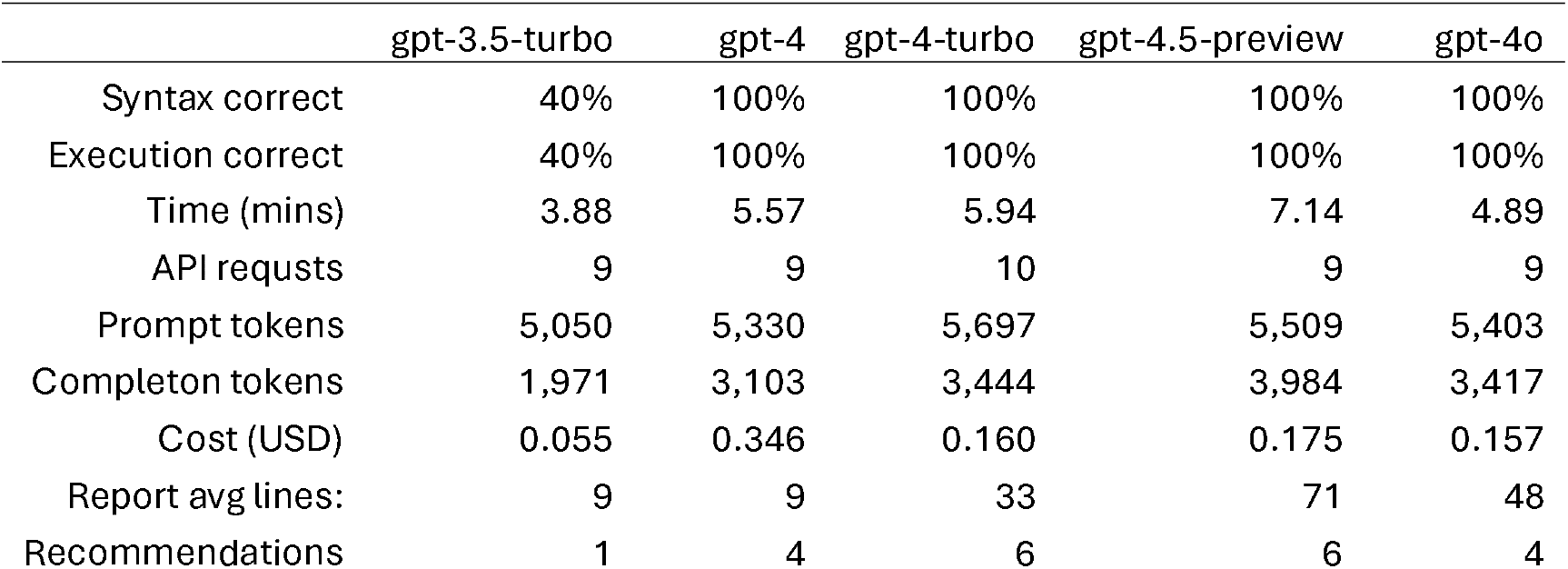
Benchmarking of different openAI models: Comparison of different OpenAI models (gpt-3.5-turbo, gpt-4, gpt-4-turbo, gpt-4.5-preview, and gpt-4o) based on key performance metrics. The models were evaluated on syntax correctness, execution correctness, response time, API request count, token usage, cost, average report length, and number of recommendations generated. Execution time is reported in minutes, API usage reflects the number of requests made per model, and token counts are split into prompt and completion tokens. Cost is reported in USD and reflects pricing as of March 2025. The results highlight differences in computational efficiency, accuracy, and cost-effectiveness across models.

### Agent-Based Reporting System

Upon workflow completion, FlowAgent leverages its natural language capabilities to interpret results and generate context-aware insights that integrate multiple analysis metrics with biological interpretations (Figure 3). Using its agents, the system produces structured reports that summarise quality control metrics, quantification performance, and resource monitoring, ensuring that bioinformatics analyses remain robust, reproducible, and interpretable while linking computational outputs to actionable biological insights. It highlights any potential issues as well as offering recommendations for enhancing future workflows and guiding downstream analyses and result interpretation. For example, it may inform the user about low alignment rates, potential batch effects, or suboptimal normalisation strategies, or suggest additional considerations when handling noisy datasets, optimising computational efficiency, or dealing with samples that are under sequenced, ensuring a more informed decision-making process for subsequent analyses.

**Figure 3:**
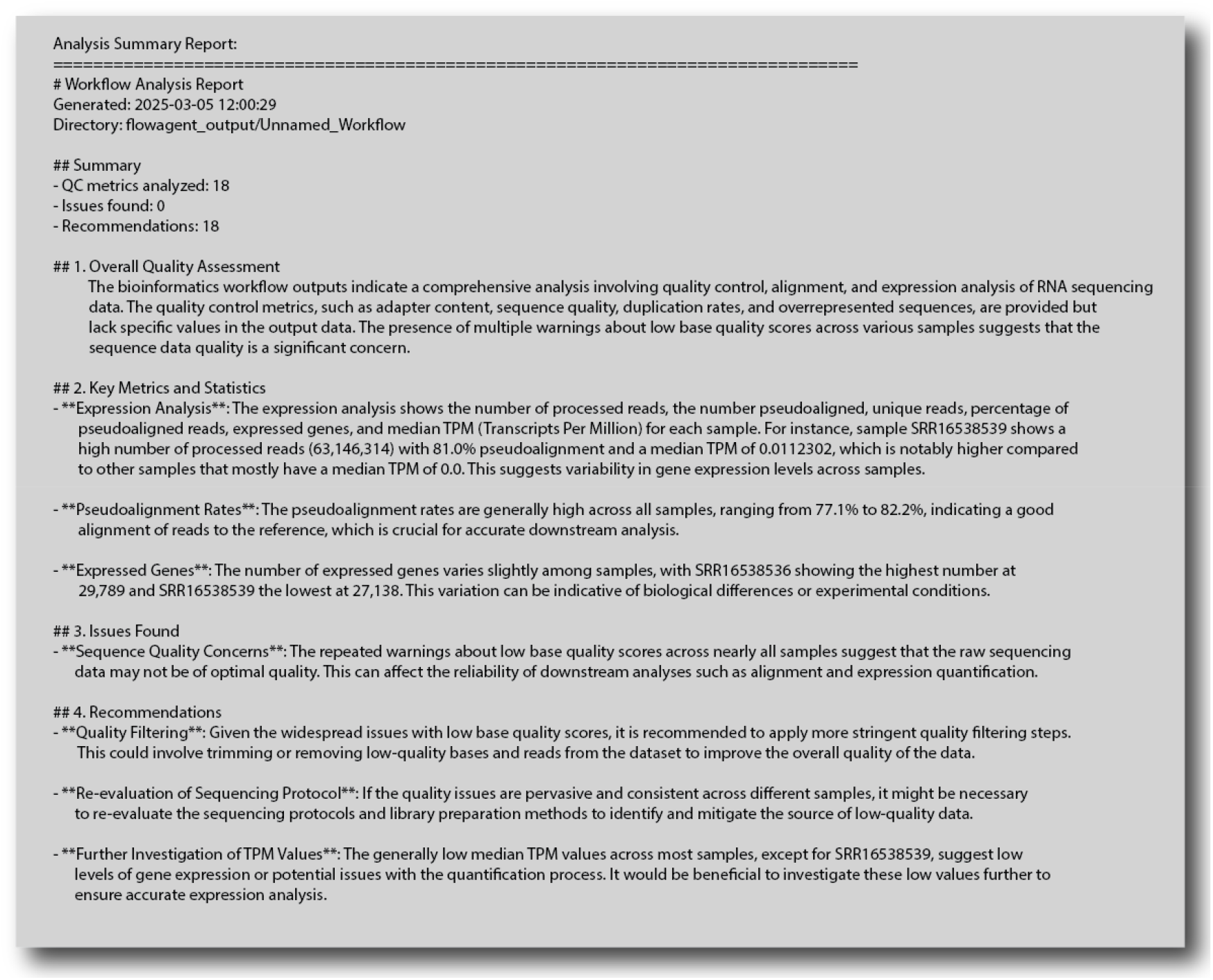
Evaluation of workflow performance post-execution by AgentFlow. FlowAgent’s agent-based reporting and evaluation system integrates quality control, quantification analysis, and technical monitoring. Specifically, the Quality Analysis Agent assesses sequencing data integrity and contamination levels, the Quantification Analysis Agent validates expression quantification, and the Technical QC Agent monitors computational efficiency, software dependencies, and pipeline performance. FlowAgent interprets and summarises workflow execution results, providing structured reports that incorporate multiple analysis metrics as well as highlighting potential issues and giving recommendations for future workflow improvements or analysis considerations. The reporting output shown is following running the gpt-4-turbo model.

## Conclusions

FlowAgent is a significant advancement in bioinformatics workflow management, addressing the limitations of traditional and emerging AI-driven systems. While conventional tools ensure structure and reproducibility, they often require specialist oversight and extensive programming expertise, creating bottlenecks. FlowAgent overcomes these challenges by integrating adaptive execution, intelligent quality control, and agent-based automation, enabling researchers—particularly those in wet-lab environments—to run complex analyses with minimal training. Its dynamic, AI-driven workflows continuously refine analyses in real time, ensuring robust, reproducible results while engaging researchers through a lab-in-the-loop approach. By using LLMs, FlowAgent simplifies workflow design through natural language, enhancing accessibility and functionality. Its modular architecture allows for seamless integration of new tools, making it scalable and adaptable to the evolving needs of bioinformatics. As computational optimisation and federated computing advance, FlowAgent will remain a powerful, intelligent solution for managing bioinformatics workflows across diverse research applications.

While the current implementation focuses primarily on upstream bioinformatics workflows, FlowAgent is expected to evolve towards fully flexible downstream analyses. Future development will prioritise customisable, modular workflows that accommodate diverse analytical endpoints, including pathway enrichment, regulatory network reconstruction, and integrative multi-omics analysis. Additionally, FlowAgent will extend its support for niche workflows and specialised assays, such as spatial transcriptomics, ensuring compatibility with emerging technologies. By incorporating adaptive decision-making and intelligent automation, FlowAgent will facilitate more sophisticated hypothesis generation, experimental design recommendations, and real-time workflow optimisation, positioning it as a comprehensive bioinformatics automation framework for diverse research applications.

